# Image Processing in DNA

**DOI:** 10.1101/2019.12.15.877290

**Authors:** Chao Pan, S. M. Hossein Tabatabaei Yazdi, S Kasra Tabatabaei, Alvaro G. Hernandez, Charles Schroeder, Olgica Milenkovic

## Abstract

The main obstacles for the practical deployment of DNA-based data storage platforms are the prohibitively high cost of synthetic DNA and the large number of errors introduced during synthesis. In particular, synthetic DNA products contain both individual oligo (fragment) symbol errors as well as missing DNA oligo errors, with rates that exceed those of modern storage systems by orders of magnitude. These errors can be corrected either through the use of a large number of redundant oligos or through cycles of writing, reading, and rewriting of information that eliminate the errors. Both approaches add to the overall storage cost and are hence undesirable. Here we propose the first method for storing quantized images in DNA that uses signal processing and machine learning techniques to deal with error and cost issues without resorting to the use of redundant oligos or rewriting. Our methods rely on decoupling the RGB channels of images, performing specialized quantization and compression on the individual color channels, and using new discoloration detection and image inpainting techniques. We demonstrate the performance of our approach experimentally on a collection of movie posters stored in DNA.

## 1. INTRODUCTION

DNA-based data storage has recently emerged as a viable alternative to classical storage devices that can be used to record bits at a nanoscale level and preserve them in a nonvolatile fashion for thousands of years [1, 2, 3, 4, 5, 6, 7, 8, 9]. Almost all existing DNA-based data recording architectures store user content in synthetic DNA strands of length 100 − 1000 base-pairs, organized within large unordered pools, and retrieve desired information via next-generation (e.g., HiSeq and MiSeq) or third-generation nanopore sequencing. Although DNA sequencing can be performed at a very low cost, de novo synthesis of DNA oligos with a predetermined content still represents a major bottleneck of the platform. Synthetic DNA platforms are prohibitively expensive compared to existing optical and magnetic media [6]. Furthermore, synthetic DNA-based storage systems have error-rates of the order of 10^−3^ that by far exceed those of existing high-density recorders. Synthesis errors include both symbol errors as well as *missing oligo errors* which are unique to this type of storage media and refer to the fact that one may not be able to cover all sub-strings of the user-defined string. Missing oligos represent serious obstacles to accurate data retrieval as they may affect more than 20% of the product. To address this type of error, Grass et al. [4] proposed using Reed-Solomon codes at both the oligo and pool of oligo level to ensure that missing strings may be reconstructed from combinations of redundantly encoded oligos. Unfortunately, adding redundant oligos further increases the cost of the system as the oligos have to be sequenced to determine the missing oligo rate in order to add the correct amount of redundancy.

We propose a new means of archiving images in DNA in which the missing and erroneous oligos are corrected through specialized learning methods, rather than expensive coding redundancy. The gist of our approach is to first aggressively quantize and compress colored images by specialized encoding methods that separately operate on the three color channels, RGB. Our quantization scheme reduces the image color pallet to 8 intensity levels per channel, and compresses intensity levels through a combination of Hilbert-space filling curves, differential and Huffman coding. Given that compression may lead to catastrophic error-propagation in the presence of missing or mismatched oligos, we also introduce very sparsely spaced markers into the oligo codes in order to resynchronize positional pixel information when this is lost. No error-correcting redundancy is added to the pool in order to further save in synthesis cost, and instead, the retrieved corrupted images are subjected to specialized image processing techniques that lead to barely distorted outputs. Our scheme combines automatic detection of discolorations in images with inpainting based on EdgeConnect [10] and smoothing via bilateral filtering [11]. We experimentally tested our proposed DNA image processing scheme on a pool of 11, 826 oligos of length 196 basepairs each, purchased from Integrated DNA Technologies (IDT).

The paper is organized as follows. Section 2 contains a description of our color image encoding scheme, while Section 3 describes the oligo synthesis, amplification, image processing procedures and experimental results.

## 2. THE ENCODING PROCEDURE

Our two-step encoding procedure first translates an image file into 24 binary strings, and then converts the binary strings into DNA oligos for storage and amplification. A detailed description of each step used in the process is provided below.

### Converting image files to binary strings

The first step in the procedure is *RGB channel separation and quantization*. First, we split the color images into three color channels, R (red) G (green) B (blue), and then perform 3-bit quantization of the values in each channel. More precisely, the image *I* is represented by a three-dimension tensor of size *m*×*n*×3, i.e., *I* ∈ [256]^*m*×*n*×3^, which we split into three matrices **R**, **G**, **B** of size *m* × *n* each. Next, we perform 3-bit quantization of each color matrix, leading to intensity values mapped from 0 − 255 to 0 − 7. More specifically, we use the following quantization rule for all three channels:

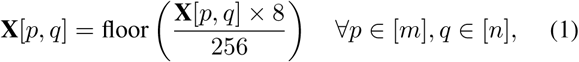

where **X** ∈ [8]^*m*×*n*^ is the quantized matrix for **X** ∈ {**R**, **G**, **B**}.

### Converting 2D images into 1D oligo strings

There exist several methods for converting a matrix into a string so as to nearly-optimally preserve 2D image distances in the 1D domain, such as the Hilbert and Peano space-filling curve. The Hilbert space-filling curve, shown in Figure 1, provides a good means to capture 2D locality [12, 13] and is the method of choice in our conversion process. Note that the Hilbert curve is standardly used on square images, so we adapt the transversal implementation to account for matrices with arbitrary dimensions (the detailed description of the mapping is postponed to the full version of the paper). After the mapping, the matrices **R**, **G**, **B** are converted into vectors *V*_*R*_, *V*_*G*_, *V*_*B*_, respectively.

**Fig. 1.**
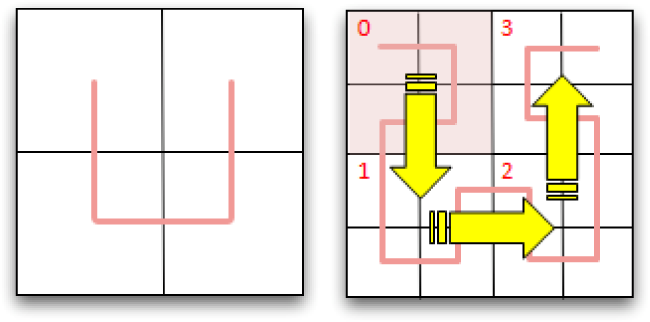
Hilbert curves for 2 × 2 and 4 × 4 squares.

### Partitioning color channels according to levels

Upon quantization, the values in *V*_*R*_, *V*_*G*_, *V*_*B*_ lie in {0, …, 7}. We next decompose each vector into strings of possibly different lengths according to the intensity value. Specifically, *V*_*R*_ is decomposed into *L*_*R*, 0, …_, *L*_*R*, 7,_ where the vector *L*_*R,j*_ contains the indices of the elements in *V*_*R*_ whose value equals *j, j* ∈ [8]; the same procedure is performed for the vectors *V*_*G*_, *V*_*B*_. An example decomposition may read as:

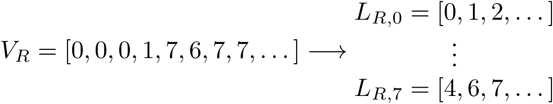

Note that the elements in *V*_*i*_ are assigned to *L i,j* in order, *i* ∈ {*R, G, B*}, 0 ≤ *j* ≤ 7. Hence each vector *L*_*i,j*_ contains increasing values, a fact that we exploit in our reconstruction procedure. Given the Hilbert scan, one may expect the differences between adjacent entries in each of the vectors *L*_*i,j*_ to be small with high probability. Therefore, splitting a vector into individual levels enables subsequent differential encoding [14]. Moreover, since the level information is split among different vectors, we will also be able to correct distortions in the images in the presence of errors. In summary, after the RGB decomposition and level partition, each image is represented by 24 vectors. **Differential encoding** converts a string into another string containing the initial value of the original and the differences between consecutive values, summarized in vectors denoted by *D*_*i,j*_. In order to prevent catastrophic error propagation, we set 3% of the values in each *D*_*i,j*_ to their original undifferentiated values and prepend to them the symbol −1. We also append an additional − 2 to each *D*_*i,j*_ to indicate the end of the vector. For example, a typical pair of *L*_*i,j*_ and *D*_*i,j*_ may be of the form:

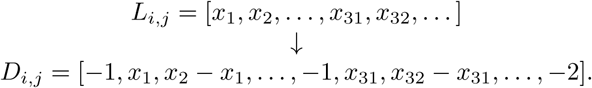

Note that as *L*_*i,j*_ has increasing values, the symbols −1 and −2 cannot be confused with information-bearing values in *D*_*i,j*_. **Huffman coding** [15] is performed after differential coding, and all values in *D*_*i,j*_ are used to construct the Huffman code dictionary. This results in a collection of binary strings *B*_*i,j*_, *i* ∈ {*R, G, B*}, 0 ≤ *j* ≤ 7.

### Conversion of binary strings into DNA oligos

The binary information is converted into oligo strings over the alphabet {A,T,G,C} of length 196 nucleotides. Each oligo contains a unique block-id for its position in the original string. If needed, some strings are padded with dummy values to ensure uniform lengths. Once again, −2 is used to indicate the end of the vector. In addition, each DNA oligo includes a prefix primer, address, an information block and suffix primer.

### Mapping binary sequences to DNA blocks

To ensure a high quality of the synthetic product, we perform constraint coding by imposing a maximum runlength-3 constraint for the symbols C and G and ensuring a GC content in the range 40 − 60% [3]. The constrained coder maps 18 and 22-bit sequences into 10 and 13 nucleotide DNA oligos, respectively. This constrained code, along with the color code, is the only source of redundancy in the encoding procedure.

### Primer sequences

We add to each DNA oligo a prefix and suffix primer, used for PCR amplification of the single stranded DNA oligos. To allow for random access, we choose 8 pairs of primers of length 20, one for each level, all of which are at a Hamming distance ≥10 nucleotides. The primers are paired up so as to have similar melting temperature, which allows for all oligos to be amplified in the same cycle.

### Address sequences

Strings of length 13 are added to the DNA oligos following the primers in order to represent the address of the information blocks contained. The first 3 nucleotides of the address encode the color (RGB). Since color information is highly important for reconstruction, we present it in redundant form as R = ‘ATC’, G = ‘TCG’, B = ‘GAT’. This allows for single-error correction in the color code. The second part of the address is of length 10 nucleotides, encoding a 18-bit binary string including the index of the corresponding image file, the index of the color level and the index of the information block within that level.

**Information blocks** are added to the oligos between the address and suffix primer, including 11 blocks of length 13 nucleotides. The total length of the information block is 143 nucelotides. Overall, with the compression scheme and additional addressing information added, 8, 654, 400 bits of the original images are converted into 2, 317, 896 nucleotides. The encoding steps are summarized in Figure 2.

**Fig. 2.**
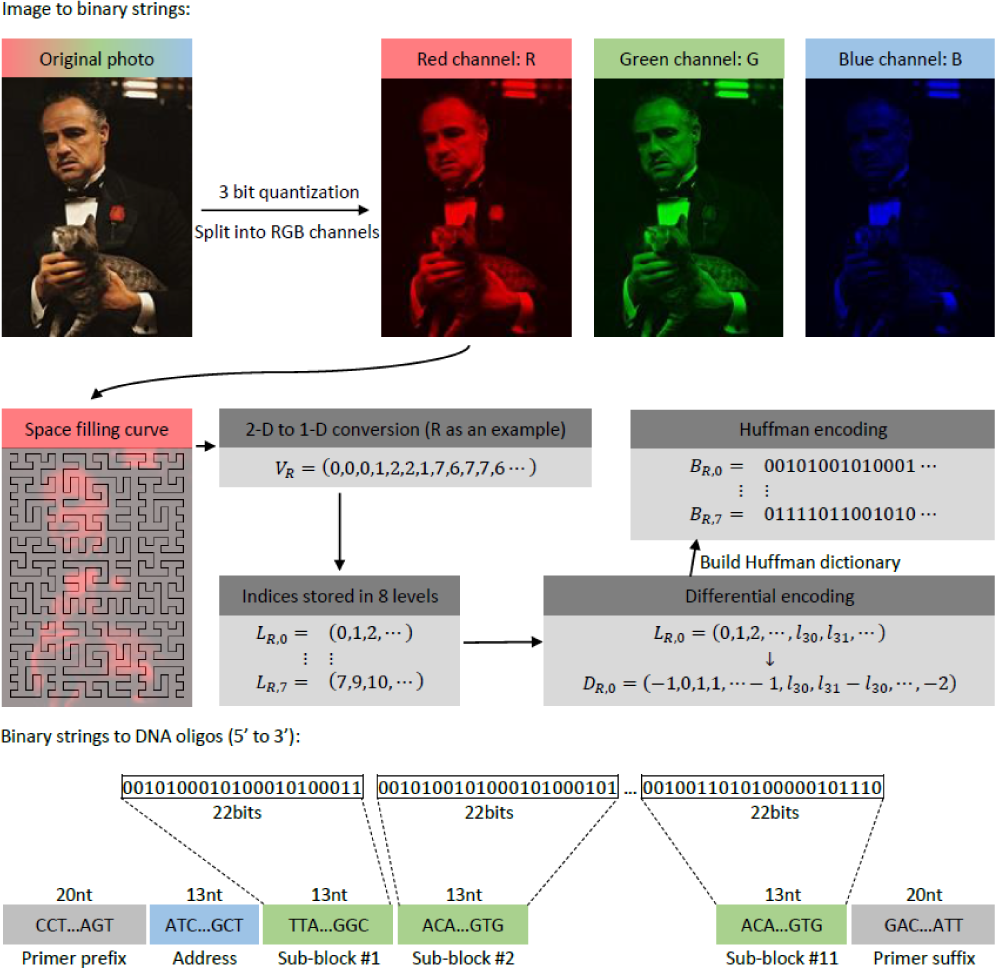
Schematic depiction of the encoding procedure.

## 3. DNA IMAGE PROCESSING AND EXPERIMENTS

The 11, 826 DNA oPools oligos were ordered from IDT (https://www.idtdna.com/pages/products/custom-dna-rna/dna-oligos/custom-dna-oligos/opools-oligo-pools). They were PCR amplified and the PCR products were then converted into a shotgun sequencing library with the Hyper Library construction kit from Kapa Biosystems (Roche). The library was quantitated by qPCR and sequenced on one ISeq flowcell for 251 cycles from one end of the fragments. The fast file was generated with the Illumina bcl2fastq v2.20 Conversion Software. As each oligo read may contain errors that arise both during synthesis and sequencing, we first *reconstructed a consensus sequence* via sequence alignment to exploit the inherent redundancy of the read process. After the whole writing, reading and consensus process, we obtained 10, 981 perfectly reconstructed oligos, 745 oligos with symbol errors that do not cause obvious defects in the reconstructed images, and 100 oligos with large corruption levels or completely missing from the pool.

The decoding procedure operates on the consensus reads and reverses the two-step encoding process. Due to lack of space, a detailed description of the procedure is omitted.

### Converting DNA consensus strings into binary strings

If some oligo unique identifiers are corrupted by errors during the synthesis or sequencing process, we replace the erroneous identifier by a unique string at smallest Hamming distance from it. Each DNA block is converted into some binary string, although this string may be wrong and cause visible discolorations in the image.

### Image processing

An example illustrating the image corruptions caused by erroneous/missing oligos is shown in Figure 3. Small blocks with the wrong color can be easily observed visually, and they are a consequence of only 10 missing oligos. To correct the discolorations automatically, we propose a three-part image processing procedure. The first step consists in detecting the locations with discolorations, masking the regions with discolorations and subsequently treating them as missing pixels. The second step involves using deep learning techniques to inpaint the missing pixels. The third step involves smoothing the image to reduce both blocking effects caused by aggressive quantization and the mismatched inpainted pixels.

**Fig. 3.**
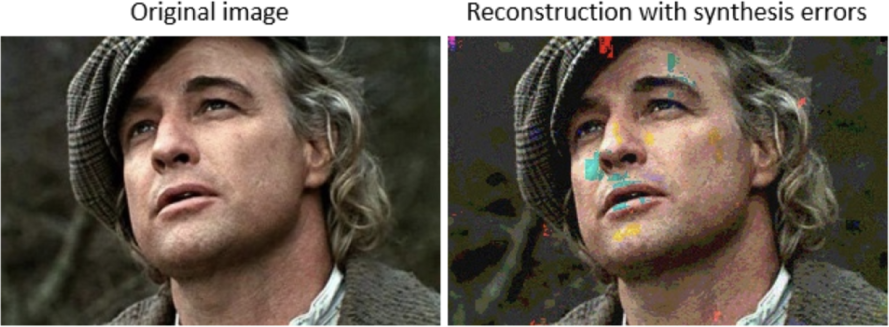
The reconstructed image with discolorations.

### Automatic discoloration detection

To the best of our knowledge, detecting arbitrarily shaped discolorations is a difficult problem in computer vision that has not been successfully addressed for classical image processing systems. This is due to the fact that discolored pixels usually have simultaneous distortions in all three color channels of possibly different degrees. However, detecting discolorations in DNA-encoded images is possible since with high probability, only one of the three color channels will be corrupted due to independent encoding of the RGB components. Figure 4 illustrates this fact, as erroneous pixels in different channels do not overlap. Within the correct color channels, pixels have neighbors of similar level, while within the erroneous channel, pixels have values that differ significantly from those of their neighbors. Figures 5 (a)(b)(c) illustrates that pixels with the smallest *t* = 15 frequencies in the difference vectors indeed correspond to almost all erroneous regions in the red channel. The results of our detection scheme are depicted in Figure 5(d)(e), for *t* = 18. Note that the whitened out regions are treated as missing data, and filled in using inpainting techniques.

**Fig. 4.**
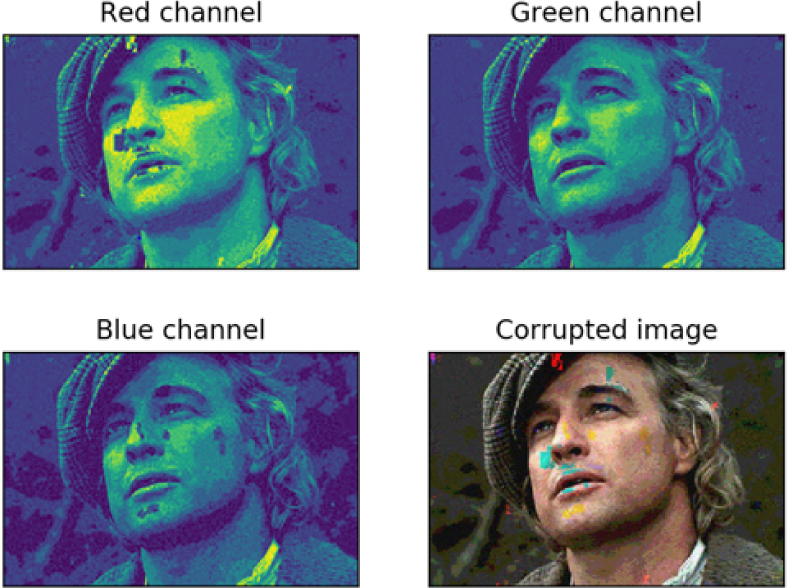
Non-overlapping errors in different color channels of the image encoded in DNA.

**Fig. 5.**
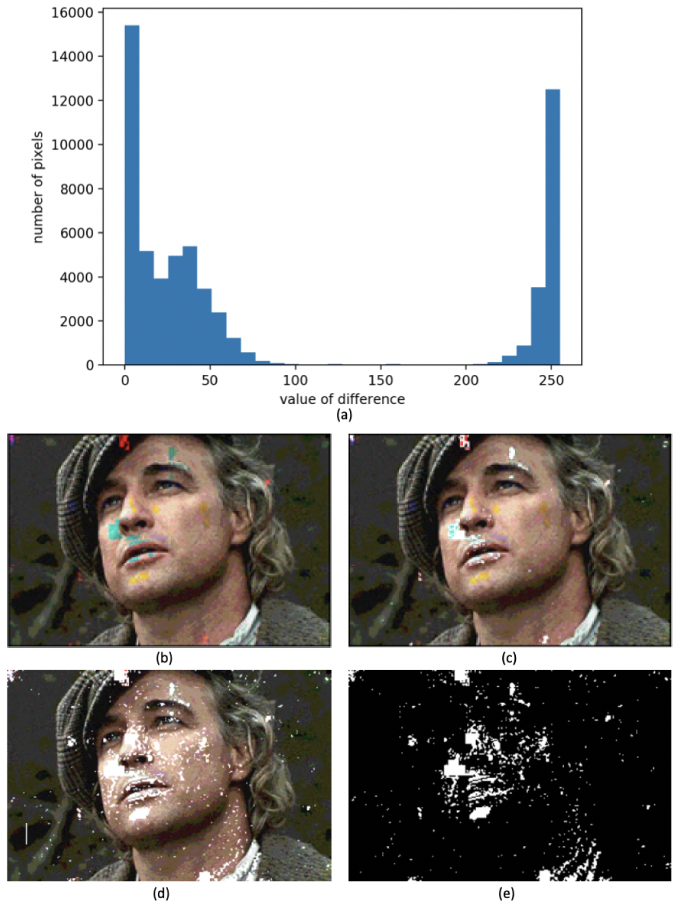
(a) Histogram of the values in the matrix **R − G**. (b) Corrupted image. (c) Discolored regions in the red channel that have been whitened out. (d) An erroneous reconstruction with masking. (e) The Image of the mask.

**Image inpainting**, or image completion, is a method for filling out missing regions in an image. There exist several methods for image inpainting currently in use, including diffusion-based, patch-based [16] and deep learning approaches [10]. The former two methods use local or non-local information only within the target image itself which leads to poor performance when trying to recover complex details in large images. On the other hand, deep-learning methods such as EdgeConnect [10] combine edges in the missing regions with color and texture information from the remainder of the image to fill in the missing pixels. Since the encoded movie posters have obvious edge structures, we inpainted the images using EdgeConnect with the result shown in Figure 6(a).

**Fig. 6.**
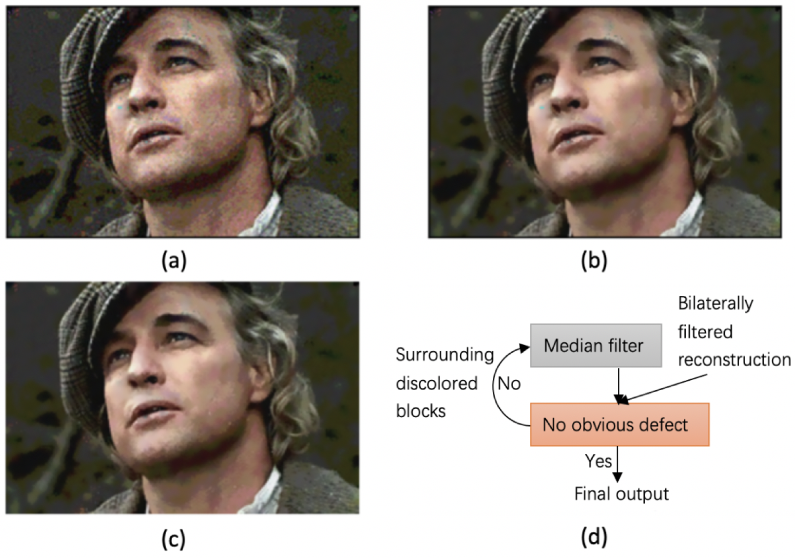
(a) Inpainting before smoothing. (b) Inpainting after smoothing. (c) Refined output of the inpainting procedure. (d) Description of the refining procedure.

### Smoothing

Although the problem of discoloration may be addressed through inpainting, the reconstructed images still suffer from mismatched inpaints and blocking effect caused by quantization. To further improve the image quality we perform smoothing through bilateral filtering [11] that tends to preserve the edges structures. The smoothing equations read as:

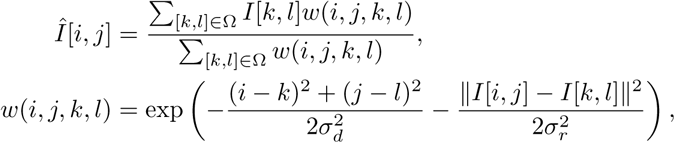

where *I* denotes the original image and *Î* the filtered image, Ω is some predefined window centered at the coordinates [*i, j*], and *σ*_*r*_ and *σ*_*d*_ are parameters that control the smoothing differences for intensities and coordinates, respectively. The filter performs Gaussian blurring on background regions but respects edge boundaries in the image. The result of smoothing with 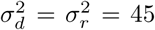 and Ω of the form of a 9 × 9 square is shown in Figure 6(b), and no obvious discolorations are detectable. Furthermore, in order to address other possible impairments, we also used the positions of error blocks obtained from the discoloration detection platfrom to perform adaptive median smoothing around erroneous regions. The output of this iterative process is illustrated in Figure 6(c)(d).

